# Caveolin-1-dependent signaling and survival in distinct Ewing sarcoma cell states

**DOI:** 10.1101/2024.09.23.614468

**Authors:** Dagan Segal, Xiaoyu Wang, Hanieh Mazloom-Farisbaf, Felix Y. Zhou, Divya Rajendran, Erin Butler, Bingying Chen, Bo-Jui Chang, Khushi Ahuja, Averi Perny, Kushal Bhatt, Dana Kim Reed, Diego H. Castrillon, Jeon Lee, Elise Jeffery, Lei Wang, Khai Nguyen, Noelle S. Williams, Stephen X. Skapek, Satwik Rajaram, Reto Fiolka, Khuloud Jaqaman, Gary Hon, James F. Amatruda, Gaudenz Danuser

## Abstract

Plasticity is a hallmark function of cancer cells, yet the mechanisms that enable dynamic switching between survival states remain incompletely understood. Here, we identify Caveolin-1, a membrane-domain scaffolding protein, as a regulator of survival signaling in a Ewing sarcoma (EwS) cell line. Single-cell analyses reveal a distinct subpopulation of EwS cells marked by high CD99 and elevated Caveolin-1 expression. These CD99^High^ cells exhibit unique morphology, transcriptional programs, and markedly enhanced survival both under chemotherapeutic challenge and *in vivo.* Importantly, CD99^High^ and CD99^Low^ states are reversible, providing EwS cells with a flexible route to survival-oriented plasticity. Mechanistically, we show that Caveolin-1 in CD99^High^ cells orchestrates PI3K/AKT survival signaling by modulating recruitment of PI3K activity to the plasma membrane. We propose that the CD99^High^ state establishes a Caveolin-1-driven signaling architecture that supports survival through mechanisms distinct from those used by CD99^Low^ cells. These findings uncover a dynamic state transition in EwS cells and position Caveolin-1 as a regulator of cell-state-dependent survival signaling.

## INTRODUCTION

Cellular adaptation is a hallmark of cancer, driving intratumoral heterogeneity and contributing to tumor growth, metastasis, and therapeutic resistance ^1^. Across many cancer types and under varying stresses, phenotypically distinct yet genetically similar subpopulations with different growth and metastatic capabilities have been identified ^2^. However, the underlying mechanisms often remain poorly understood. This gap in knowledge is especially critical in pediatric cancers, which tend to exhibit a low mutational burden beyond a single driver, despite variability in patient outcome ^3, 4^. In this work, we focus on Ewing sarcoma (EwS) as a prototype of pediatric cancer. EwS is the second most common malignant bone tumor occurring in children and young adults, characterized by a single chromosomal translocation leading to the formation of a chimeric fusion oncogene transcription factor, most commonly EWSR1::FLI1 ^5, 6^. While much research has focused on EWSR1::FLI1 as the putative driver of distinct EwS cell states ^7, 8^, more recent works have identified EwS cell states with distinct gene expression signatures not directly associated with EWSR1::FLI1-mediated transcriptional regulation ^9–11^. In either case, the signaling mechanisms that drive segregation of cell states remain largely unknown.

One possible mechanism driving distinct cell states could be through regulation of signaling pathways on the plasma membrane, which integrates signals from the extracellular environment through targeted, localized signaling cascades. Differential organization of modified lipids or activated receptors on the cell surface into subcellular domains can affect downstream signaling^12, 13^. A prominent example of such domains can be observed in caveolae, cave-like membrane invaginations that are rich in cholesterol and sphingolipids^14^ and have roles in signal transduction ^15–20^. Caveolae are considered specialized lipid domains, with effects on membrane fluidity and lipid species composition^21, 22^. These structures are prominently found in a subset of cell types, such as adipocytes and muscle cells, and have been implicated in many cancers^23, 24^, including Ewing Sarcoma ^25, 26^. Several studies have demonstrated that perturbation of caveolae through mechanical means ^27–29^ or perturbations of the key structural protein Caveolin-1 affect canonical cancer signaling pathways, including Src^16, 19, 20^, Erk1/2^26, 30^, and PI3K/Akt activation ^31–33^, which has previously been linked to Cav-1 mediated cell survival^34^. Some of these studies have demonstrated differential subcellular organization of specific signaling molecules upon loss of caveolae ^30, 35^. However, whether and how caveolae-based organization of signaling contributes to cellular plasticity in development and disease remains unclear.

In this study, we identified a distinct cell population in an EwS cell line characterized by reduced proliferation but enhanced viability both *in vivo* and in response to chemotherapeutic treatment. We refer to this population as CD99^Hi^ cells, defined by their elevated expression of the surface marker CD99. CD99^Hi^ cells display high endogenous Caveolin-1 levels, are enriched in caveolae, and depend on Caveolin-1 for their survival advantage. Mechanistically, we show that loss of Caveolin-1 prevents CD99^Hi^ cells from recruiting active PI3K to the plasma membrane, thereby impairing activation of the PI3K/AKT survival pathway. Together, these findings reveal a mechanism by which EwS cells can adapt survival signaling in a context-dependent manner.

## RESULTS

### High CD99 marks a distinct Ewing Sarcoma cell state

Nongenetic heterogeneity in cancer cells has increasingly been recognized as a critical driver of aggressive disease progression and treatment resistance ^1, 2^. To study heterogeneity in EwS cell populations, we first used single cell transcriptomics to identify cell surface markers that would readily enable isolation of distinct subpopulations for further downstream characterization (Figure 1A). Given that cell surface receptors are sensitive to cleavage through standard enzymatic passaging techniques and inspired by classic studies in mesenchymal stem cells ^36–38^, we reasoned that cells passaged through mechanical means rather than enzymatic cleavage would enable the discovery of distinct subpopulations of cells, as well as preserve surface receptors intact as targets for flow cytometry. Applied to the EwS cell line TC71, single-cell transcriptomic analysis uncovered a cluster of cells with a unique transcriptional profile, characterized by enrichment of extracellular matrix (ECM) structural constituents (Figure S1A,B).

**Figure 1:**
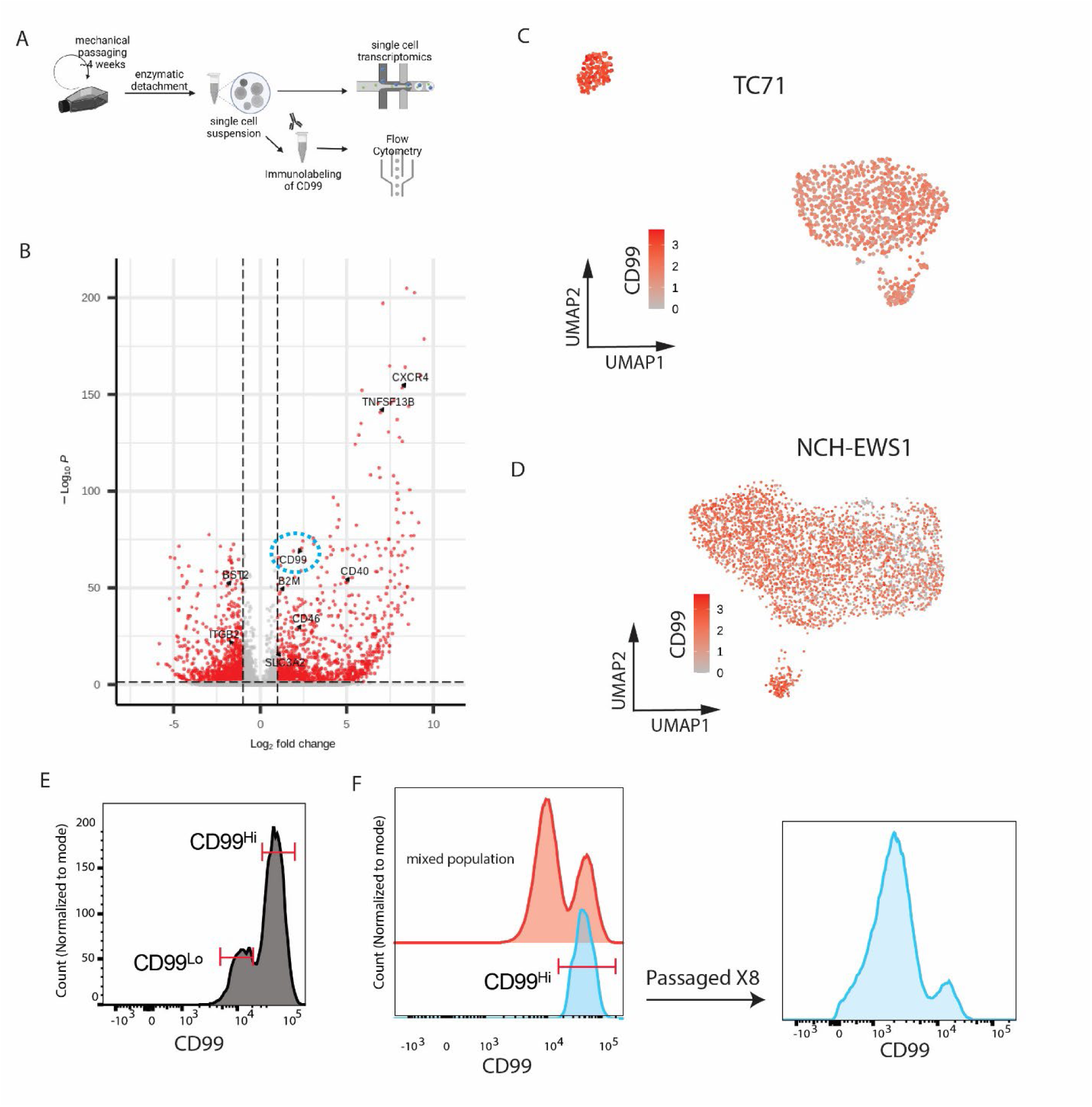
High levels of CD99 gene and protein expression defines a distinct Ewing Sarcoma cell state. **(A)** Experimental workflow. After 6-10 cycles of mechanical passaging to preserve surface receptors, cells are enzymatically detached from the dish and prepared in a cell suspension for single cell RNA seq (scRNA-seq) to profile receptor gene expression, or immunolabeling of surface-exposed receptors for flow cytometry. **(B)** Volcano plot depicting differential enrichment of transcript expression in highlighted cluster (see cluster 2 in Figure S1A) compared to all other cells. Differentially expressed cell surface receptors are indicated; CD99 is highlighted (dashed circle). **(C-D)** UMAP visualization with log normalized expression of the CD99 gene in individual cells color-coded in TC71 cells **(C)**, or in NCH-EWS1 patient-derived xenograft (PDX) cells **(D)**. **(E)** Flow cytometry histogram depicting relative intensity of cell surface exposed CD99 as measured by immunolabeling. The histogram displays a bimodal distribution of *CD99 High* (CD99^Hi^) and *CD99 Low* (CD99^Lo^) cell populations. Distinct populations are highlighted by red bars. **(F)** CD99^Hi^ cells (blue, left) were isolated from a mixed population of CD99^Lo^ and CD99^Hi^ cells (red). The isolated CD99^Hi^ cells were reevaluated by flow cytometry (blue, right) after expansion and several rounds of passaging. The recurrent bimodal distribution of CD99 expression suggests that CD99^Hi^ cells can shift between CD99^Hi^ and CD99^Lo^ cell states.

We profiled the relative expression of transcripts of surface receptors between these two populations. Of 348 previously validated targets^39^, seven surface receptors were enriched in the distinct cluster (Figure 1B). Among those was the glycoprotein CD99 (Figure 1B,C), a commonly used biomarker for EwS^40^. Based on their distinct CD99 expression levels, we designate these populations as CD99^Hi^ and CD99^Lo^ cells. A CD99^Hi^ cluster was also similarly observed in NCH-EWS1, a patient derived xenograft (PDX) line (Figure 1D). The NCH-EWS1 CD99^Hi^ cluster was similarly enriched for ECM structural constituents (Figure S1B), suggesting conservation of the CD99^Hi^ cellular phenotype in EwS.

We next sought to leverage CD99 as a flow cytometry target, to isolate and further characterize the two cell populations identified. Through flow cytometric analysis in TC71 cells, we uncovered a bimodal distribution of CD99 expression, corresponding to CD99^Lo^ and CD99^Hi^ cell populations (Fig 1E). Subsequent bulk transcriptomic analysis of sorted populations revealed that TC71 CD99^Hi^ cells were enriched for gene ontology terms related to neural differentiation compared to CD99^Lo^ cells (Figure S1C). Importantly, while all EwS cells remained broadly CD99-positive by flow cytometry, mechanical passaging revealed a distinct CD99^Hi^ subpopulation in addition to a CD99^Lo^ population that overlapped with enzymatically passaged cells (Figure S1D). In contrast, the CD99^Hi^ population was lost following conventional enzymatic dissociation, suggesting that this specific high-expression state may have been overlooked in previous studies due to its sensitivity to standard dissociation procedures, despite CD99 being a canonical EwS marker. Notably, the CD99^Hi^ state appeared reversible, as isolated CD99^Hi^ cells re-established a bimodal CD99 expression pattern after several passages in culture (Fig. 1F).

### CD99^Hi^ cells display differences in cell morphology gene expression, and survival *in situ*

We subsequently utilized CD99 as a flow cytometry marker to isolate and characterize the two cell states. Strikingly, the two subpopulations exhibited distinct morphological phenotypes (Figure 2A). Imaging of TC71 cells expressing the cytoskeletal marker F-Tractin revealed that CD99^Hi^ cells displayed a flattened morphology with prominent actin bundles, whereas CD99^Lo^ cells were more rounded, formed clusters, and were enriched in filopodial protrusions (Figure 2B).

**Figure 2:**
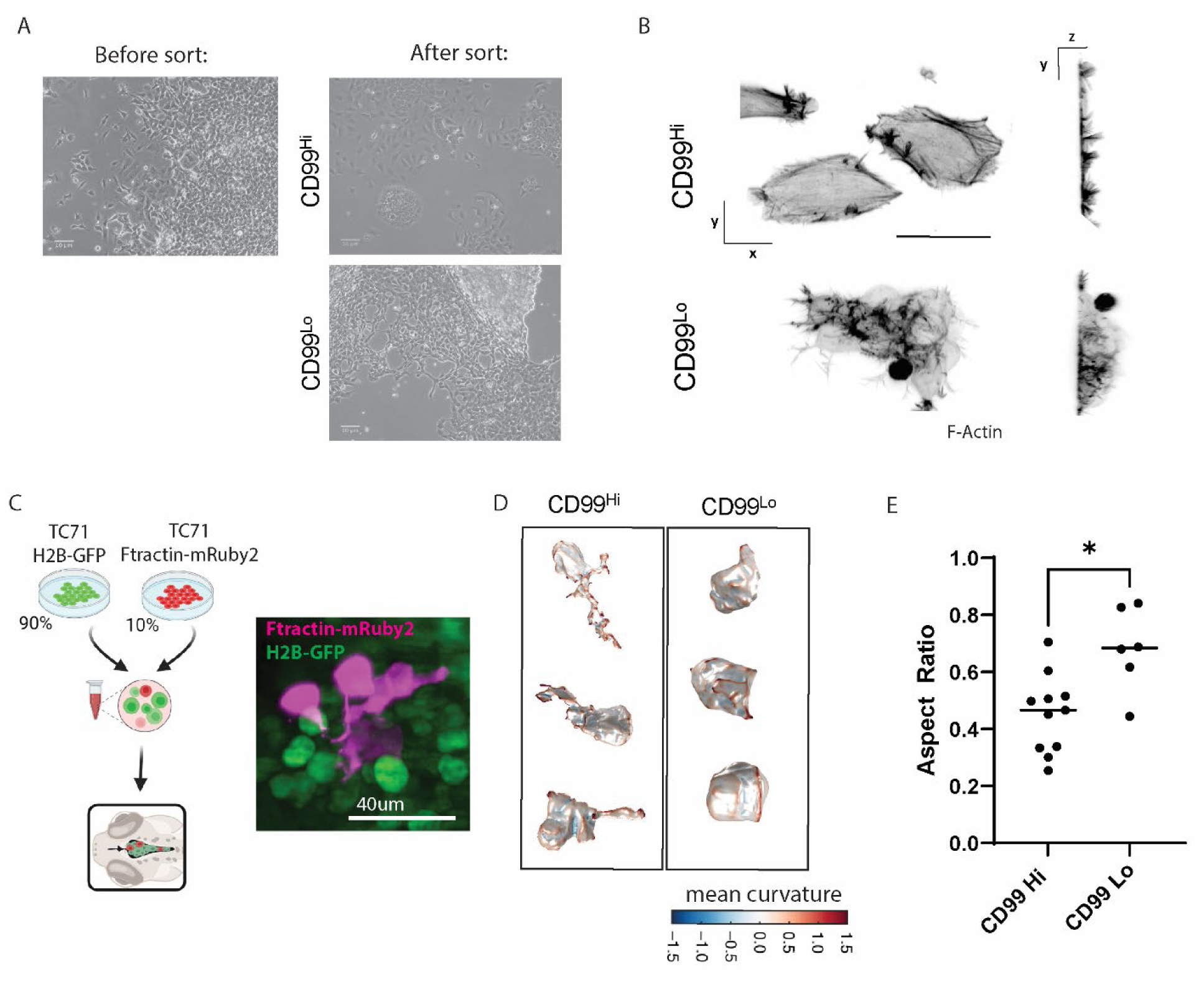
CD99^Hi^ and CD99^Lo^ cells display differences in morphology *in vitro* and in zebrafish xenografts. **(A)** Brightfield images showing TC71 CD99^Hi^ or CD99^Lo^ cell morphologies observed together before, or separately after, sorting by flow cytometry, respectively. **(B)** Maximum Intensity Projections in XY view (left) or YZ view (right) of CD99^Hi^ and CD99^Lo^ TC71 cells expressing F-Tractin-GFP. Cells were imaged in 3D using oblique plane microscopy in adherent culture. The two classes display distinct cell morphologies (scale bar = 20um). **(C) (left)** Experimental set-up to identify single-cell morphologies in xenografted cells within the hindbrain ventricle (HBV) of zebrafish larvae. In order to distinguish single-cell morphologies of individual cells, we prepared a mosaic cell mixture consisting of 90% of TC71 unsorted cells expressing the fluorescently-tagged nuclear marker H2B-GFP, and 10% of TC71 CD99^Hi^ or CD99^Lo^ cells expressing F-Tractin-mRuby2. **(right)** Maximum Intensity Projection of TC71 cells within the HBV expressing F-Tractin-mRuby2 (magenta) or H2B-GFP (green). **(D)** Volume renderings of 3 representative single cells from each class, pseudocolored by local membrane curvature. Note the highly elongated and protrusive shapes observed in CD99^Hi^ cells. **(E)** Differences in aspect ratio of individual cells, as a metric of differences in cell shape in CD99^Hi^ (n = 11 cells, 9 fish) and CD99^Lo^ cells (n=6 cells, 3 fish, * p-value<0.05, t-test). Larger aspect ratio indicates more elongated morphology.

To test whether morphological single-cell level differences were maintained *in vivo,* we utilized a previously established experimental and analytical pipeline to assay cell morphology in cancer cell xenografts in the hindbrain ventricle (HBV) of 2-day old zebrafish^41^ (Figure 2C). For this purpose, we injected a cell mixture consisting of 90% unsorted TC71 cells expressing the nuclear marker H2B-GFP, and 10% CD99^Hi^ or CD99^Lo^ cells expressing the actin tag Ftractin-mRuby2. Following fixation in 4% paraformaldehyde (PFA) and 3D imaging of the zebrafish larvae at 1-day post-injection, we computationally extracted the cell volumes of individual CD99^Hi^ or CD99^Lo^ cells for morphotype analysis, as previously described^41^. CD99^Hi^ cells displayed distinctly more irregular and elongated shapes, with prominent protrusions, than CD99^Lo^ cells (Figure 2D-E), suggesting that these EwS cells maintain the morphological differences *in vivo*.

### CD99^Hi^ cells display differences in cell survival *in vivo* and in response to chemotherapeutic drugs

We next examined broader behaviors of the CD99^Hi^ population. Growth and viability assays showed no differences between CD99^Hi^ and CD99^Lo^ cells when grown in adherent flasks (Fig S2A-B). However, CD99^Hi^ cells displayed decreased growth capabilities in soft environments *in vitro* (Fig S2C). To test whether CD99^Hi^ cells show tumor growth or survival differences *in vivo,* we performed intratibial xenografts of isolated Luciferase-expressing CD99^Hi^ or CD99^Lo^ TC71 cells, respectively, in immunocompromised mice (Figure 3A). CD99^Hi^ cells showed dramatically lower tumor burden compared to CD99^Lo^ cells (Figure 3A, Figure S2D). A computerized tomography scan also revealed that bony destruction, a phenomenon frequently found in EwS tumors and described as moth-eaten appearance of the bone^30^, was decreased in xenograft performed with CD99^Hi^ cells (Figure 3B, S2E).

**Figure 3:**
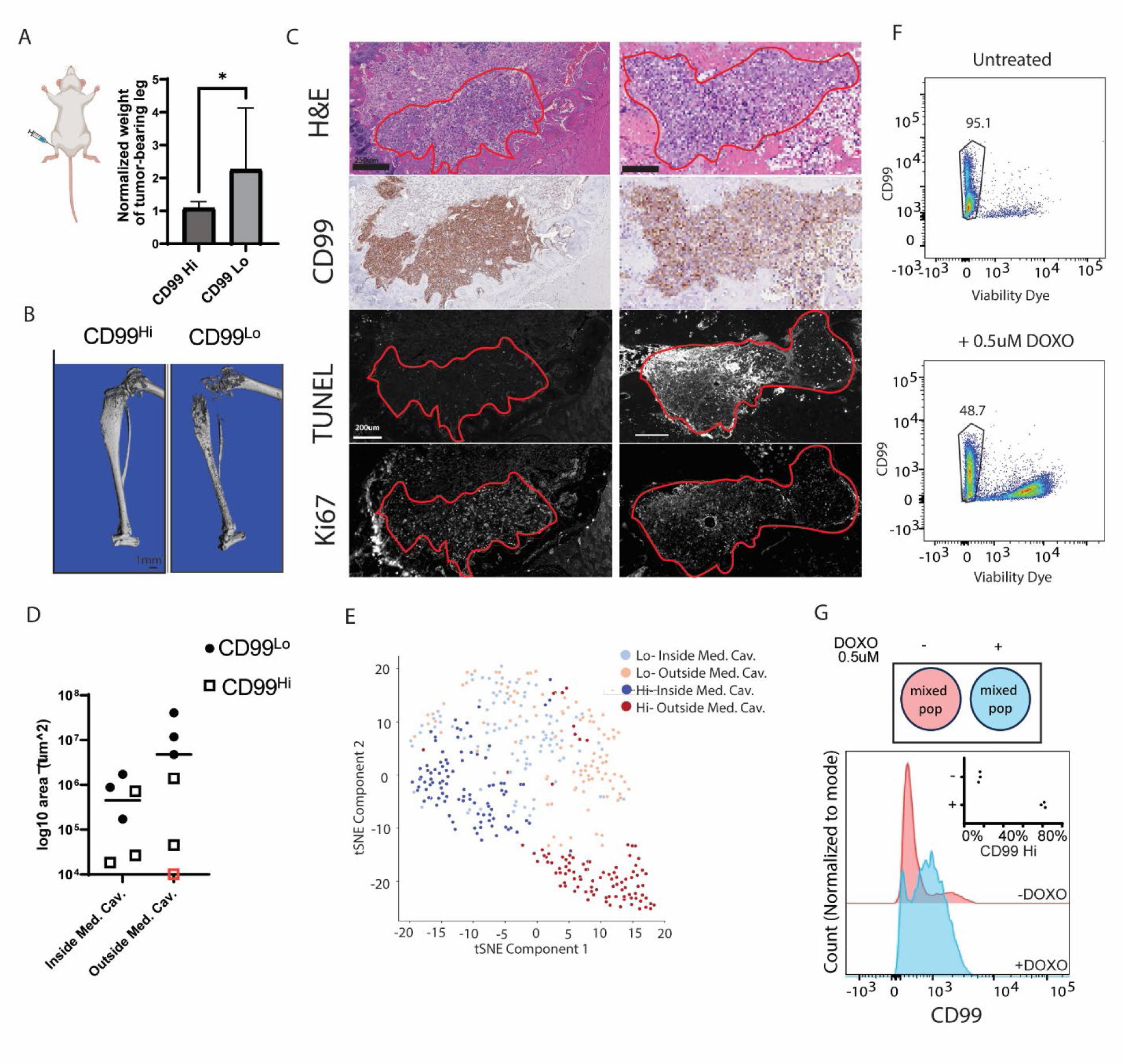
CD99^Hi^ and CD99^Lo^ cells display differences in morphology, gene expression, and survival. **(A)** (left) Schematic depicting intratibial xenografts of CD99^Hi^ or CD99^Lo^ TC71 cells into immunocompromised mice. (right) Weight of tumor-bearing leg normalized to the weight of non-tumor bearing leg of the same mouse in CD99^Hi^ (n=6) or CD99^Lo^ (n=6) xenografts (* p<0.05, t-test). **(B)** Representative computer tomography scan of tumor bearing legs depicting less bony destruction caused by CD99^Hi^ cells compared to CD99^Lo^ cells. **(C)** Histological sections of intratibial xenografts in immunocompromised mice, showing tumor-bearing areas (outlined in red) within the medullary cavity (Med. Cav.) of either CD99^Hi^ or CD99^Lo^ cells, labeled with H&E, chemically immunostained for CD99, and fluorescently stained in serial sections for the apoptotic marker TUNEL and the proliferation marker Ki67. Note the decreased TUNEL staining observed in the CD99^Hi^ tumor compared to CD99^Lo^ tumor. **(D)** Area inside the Med. Cav. of the bone, or outside of the Med. Cav., such as within the surrounding muscle. Note that while CD99^Lo^ cells have larger clusters outside the Med. Cav., CD99^Hi^ cells have comparably smaller clusters in both areas (n = 3 mice). Red square indicates CD99^Hi^ mouse with no tumor-bearing area found outside Med. Cav. Bars, mean value. (**E)** tSNE visualization of shape feature analysis on histological H&E-stained sections of intratibial xenografts of CD99^Hi^ or CD99^Lo^ cells displays differences in cell morphology between the two experimental groups. Note that while CD99^Lo^ cells show large overlap in shape feature classification regardless of seeding site, CD99^Hi^ cells show a tissue specific specialization of cell morphology. **(F)** Flow cytometry of a mixed population of CD99^Hi^ and CD99^Lo^ cells stained with Live/Dead viability dye and anti-CD99 antibody, with or without Doxorubicin treatment (DOXO, 0.5uM, 48hrs). Viable cells (Viability dye negative) constituted 95.1% of the untreated population and 48.7% of the DOXO-treated population. **(G)** Histograms depicting outlined regions from the dot plot in (F), which mark viable, CD99+ cells and show relative distributions of CD99^Lo^ and CD99^Hi^ populations grown with (+, blue) or without (-, magenta) DOXO (n=3 independent experiments). Inset, percentage enrichment of CD99^Hi^ cells in each group. CD99^Hi^ cells are systematically enriched upon drug treatment.

To study these apparent differences in tumor growth more deeply we turned to an analysis of histological sections (Figure 3C). Tumor cells were identified morphologically through H&E staining and by CD99 immunohistochemistry. Differences in CD99 expression appeared to be maintained between CD99^Hi^ and CD99^Lo^ tumors (Figure 3C, Figure S2F), suggesting distinct phenotypic cell states were sustained *in situ.* Strikingly, most tumor growth of CD99^Lo^ cells occurs through the local invasion of surrounding muscle, while both cell populations grow masses of limited size within the medullary cavity of the bone (Figure 3D, S2F). To test for survival and proliferation differences, we combined the TUNEL assay with immunostaining for the proliferation marker Ki67 in serial histological slides. For clusters of comparable size within the medullary cavity, the CD99^Lo^ cells show comparable cell proliferation but strikingly more cell death compared to CD99^Hi^ cells (Figure 3C). Differences in cell death appear to hold true for local muscle invasions as well (Figure S2G). To determine whether morphological differences were also preserved *in situ*, we performed morphological feature classification on tumor nuclei in H&E-stained histological sections. CD99^Hi^ and CD99^Lo^ cells occupied distinct regions in shape feature space (Figure 3E). Notably, CD99^Hi^ cells exhibited site-specific differences in shape features, whereas CD99^Lo^ cells showed considerable overlap in feature space across seeding sites, suggesting that CD99^Hi^ cells may have a greater capacity to adapt to diverse microenvironments.

We wondered whether the putative adaptivity of CD99^Hi^ cells also confers greater resilience to drug challenges. To test this, we grew a mixed population of CD99^Hi^ and CD99^Lo^ cells *in vitro* in the presence or absence of Doxorubicin (DOXO, 0.5uM), which serves as part of frontline therapy of EwS^40, 42^. By flow cytometry we found that DOXO treatment causes higher cell death in the CD99^Lo^ cell population (Figure 3F). Closer examination of the surviving cells revealed a significant enrichment of CD99^Hi^ cells upon drug treatment, suggesting these cells have a survival advantage under these stress conditions (Figure 3G). Taken together, the results presented thus far suggest that the CD99^Hi^ population acquires a transcriptionally and functionally distinct cell state, potentially with survival advantages in certain tissues and under cytotoxic therapy.

### Caveolin-1 emerges as a molecular signature of CD99^Hi^ State

Next, we sought to identify the potential molecular mediators that would drive functionally distinct states. Given that expression levels of the oncogene EWSR1::FLI1 have previously been implicated as a major regulator of plasticity in Ewing Sarcoma^7, 8^ and CD99 itself is a transcriptional target of EWSR1::FLI1^43^, we first tested whether differences in oncogene levels differed in the CD99^Hi^ state. While differences in CD99 expression between CD99^Hi^ and CD99^Lo^ cells were confirmed by western blot, EWSR1::FLI1 protein expression levels remained equal between the two groups (Figure S3A). We therefore sought to look more broadly to identify potential molecular signatures that could drive the functionally distinct states of CD99^Hi^ and CD99^Lo^ cells. Several initial lines of evidence converged around the scaffolding protein Caveolin-1 (Cav-1) as a potential regulator of CD99^Hi^ cells. First, Cav-1 expression was highly enriched at the transcript level in the TC71 CD99^Hi^ cluster by single-cell transcriptomics (Figure 4A). Second, through a functional proteomics screen based on a Reverse Phase Protein Array (RPPA), we observed Cav-1 as the most differentially enriched protein in TC71 CD99^Hi^ cells compared to CD99^Lo^ cells (Figure 4B). Third, ultrastructural analysis by electron microscopy revealed multiple morphological differences between the populations. Besides altered ribosomal content and endoplasmic reticulum morphology, CD99^Hi^ cells but not CD99^Lo^ cells displayed an abundance of caveolae, Cav-1-driven cholesterol-rich membrane invaginations^14^ (Figure 4C). Similarly, colocalization analysis of Cav-1 with the structural caveolar component PTRF/Cavin-1^44^ through immunofluorescence revealed high colocalization in CD99^Hi^ but not in CD99^Lo^ cells, further validating the enrichment of caveolae in CD99^Hi^ cells (Figure S3B-C). Quantitative analysis of diffusion dynamics of Cav-1 puncta showed slower diffusion in CD99^Hi^ cells (Figure S3D), another signature of stable caveolar structures^45^. Cav-1 is upregulated in metastases across several cancer types^46^, including EwS^25, 26^, and has been implicated in diverse processes, including cell signaling, membrane trafficking, and stress response, making it a promising candidate for further investigation.

**Figure 4:**
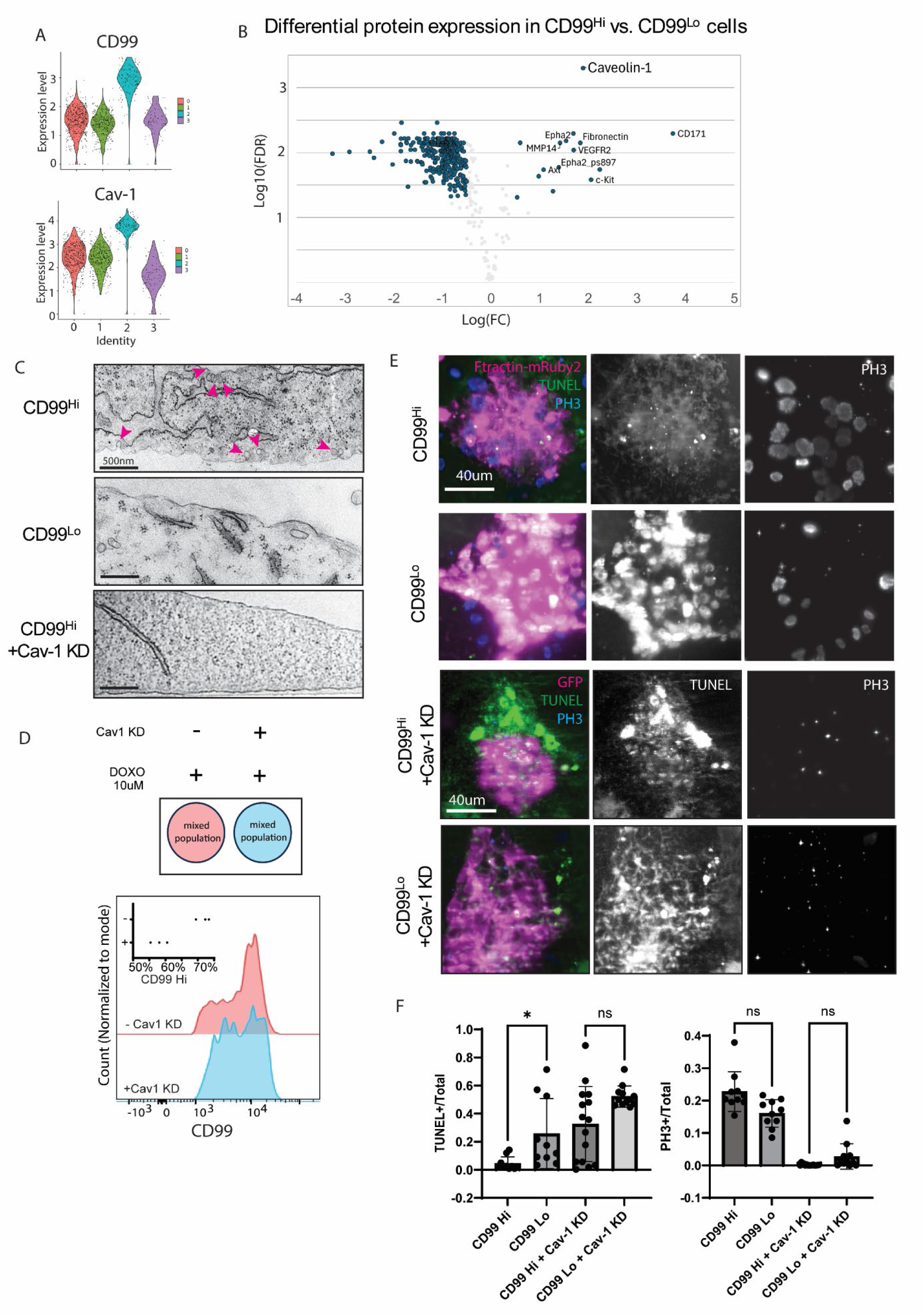
Caveolin-1 emerges as a potential driver of functions associated with the CD99^Hi^ State. **(A)** RNA expression of CD99 and Cav1 in different clusters, analyzed from scRNA-seq data (see Fig S1). **(B)** Volcano plot of differential protein expression between CD99^Hi^ vs. CD99^Lo^ cells, analyzed by Reverse Phase Protein Array (RPPA). **(C)** Representative scanning electron microscope images of TC71 CD99^Hi^, CD99^Lo^, and CD99^Hi^ cells expressing inducible Cav-1 shRNA (Cav1 KD). Magenta arrowheads highlight observed caveolae in CD99^Hi^ cells. **(D)** (Left) Experimental setup for a flow cytometry assay of a mixture of TC71 CD99^Hi^ and CD99^Lo^ cells, with (blue) or without (red) knockdown of Cav-1, grown with Doxorubicin (DOXO, 10uM, 48hrs). (Right) Histogram of CD99^Hi^ and CD99^Lo^ populations. Representative of n=3 experiments. Inset-percentage enrichment of CD99^Hi^ cells in each group (n=3). Note that enrichment of CD99^Hi^ peak under DOXO treatment is lost upon Cav1 KD. **(E)** TC71 CD99^Hi^ or CD99^Lo^ cells expressing F-tractin-mRuby2 (magenta), with (n =14 fish, n= 12 fish, respectively) or without (n = 10 fish) induced Cav1 KD were injected into the HBV of zebrafish larvae. Representative Maximum Intensity Projection images of xenografted larvae stained for apoptotic marker TUNEL (green (left), grey (middle)) and proliferative marker Phospho-histone H3 (PH3, blue (left), grey (right)) at 48hrs post-injection. **(F)** Quantification of the proportion of xenograft area positive for TUNEL (left) or PH3 (right) signal overlapping with F-Tractin-mRuby2 expressing tumor cells. Among cells with endogenous Cav-1 expression, CD99^Hi^ cells exhibited reduced cell death compared to CD99 Lo cells. Cav-1 knockdown significantly increased cell death and reduced proliferation in both CD99^Hi^ and CD99^Lo^ cells, eliminating the difference in cell death between the two populations.

To investigate whether Cav-1 drives the CD99^Hi^ state, we established a tetracycline (TET)-inducible shRNA knockdown system targeting Cav-1, combined with an RFP reporter, in TC71 cells. Strikingly, Cav-1 knockdown markedly reduced the density of membrane invaginations in CD99^Hi^ cells, demonstrating that changes in Cav-1 expression directly regulate the formation of caveolae (Figures 4C). However, inducible Cav-1 knockdown had no effect on CD99 expression (Figure S3E). Using a similar approach to target CD99 expression, we found that CD99 knockdown increased Cav-1 expression in CD99^Lo^ cells but had no effect in CD99^Hi^ cells (Figure S3F). These findings suggest that CD99 can function as an upstream regulator of Cav-1, but this regulatory relationship is bypassed once cells adopt the CD99^Hi^ state.

To determine whether the enhanced survival of CD99^Hi^ cells following DOXO treatment is Cav-1 dependent, we assessed the viability of mixed CD99^Hi^*/* CD99^Lo^ cell population in response to acute treatment with 10 μM DOXO, with or without Cav-1 knockdown. While Cav-1 knockdown did not confer an overall survival advantage under drug treatment (Figure S3G), detailed analysis of the surviving CD99+ population revealed a critical dependency on Cav-1. In cells with endogenous Cav-1 expression, DOXO treatment enriched for CD99^Hi^ cells (Figures 3G, 4D). However, this enrichment was abolished following Cav-1 knockdown, resulting in comparable proportions of CD99^Hi^ and CD99^Lo^ cells (Figure 4D). These findings demonstrate that Cav-1 is required for the survival advantage of CD99^Hi^ cells under stress from chemotherapy, suggesting a Cav-1-dependent survival signaling pathway in this cellular state.

Since CD99^Hi^ cells exhibit a survival advantage *in vivo*, we hypothesized that Cav-1 may contribute to this functional difference. Testing inducible Cav-1 knockdown in the intratibial mouse xenograft model posed substantial technical challenges due to limited tetracycline delivery to the bone microenvironment and difficulty detecting the relatively small CD99^Hi^ population in situ. We therefore turned to the zebrafish HBV xenograft model, which enables higher-resolution imaging and more accessible drug delivery. Through injection of CD99^Hi^ or CD99^Lo^ cells expressing F-Tractin-mRuby2 into the zebrafish HBV followed by chemical fixation at 2 days post-injection and apoptotic TUNEL staining, we observed significantly reduced cell death in CD99^Hi^ cells compared to CD99^Lo^ cells with endogenous Cav-1 expression (Figure 4E-F), akin to the situation in the tibial xenografts in mice. Cav-1 knockdown significantly increased cell death and reduced proliferation in both CD99^Hi^ and CD99^Lo^ cells, eliminating the difference in cell death between the two populations. These findings support a role for Cav-1 in promoting the enhanced survival of CD99^Hi^ cells *in vivo*.

### Caveolin-1 regulates PI3K/AKT signaling in CD99^Hi^ Cells

The change in viability upon drug treatment prompted us to examine changes in survival signaling pathways. By RPPA we noted in CD99^Hi^ cells an enrichment of Receptor Tyrosine Kinases that have been associated with the activation of the phosphatidylinositol 3-kinase (PI3K)/Akt survival signaling pathway, namely EphA2, VEGFR, c-Kit, and Axl (Figure 4B, Table S1)^47–50^. Indeed, western blot analysis confirmed increased active Akt at baseline in CD99^Hi^ TC71 cells (Figure 5A) and two other Ewing Sarcoma cell lines (Figure S4A) when compared to their CD99^Lo^ counter parts. Knockdown of Cav-1 reduced phosphorylated Akt levels in CD99^Hi^ cells in TC71 cells (Figure 5A), consistent with prior reports linking Caveolin-1 to Akt signaling in other cancer models^32, 34^, and indicating that a similar signaling relationship exists in EwS cells.

**Figure 5:**
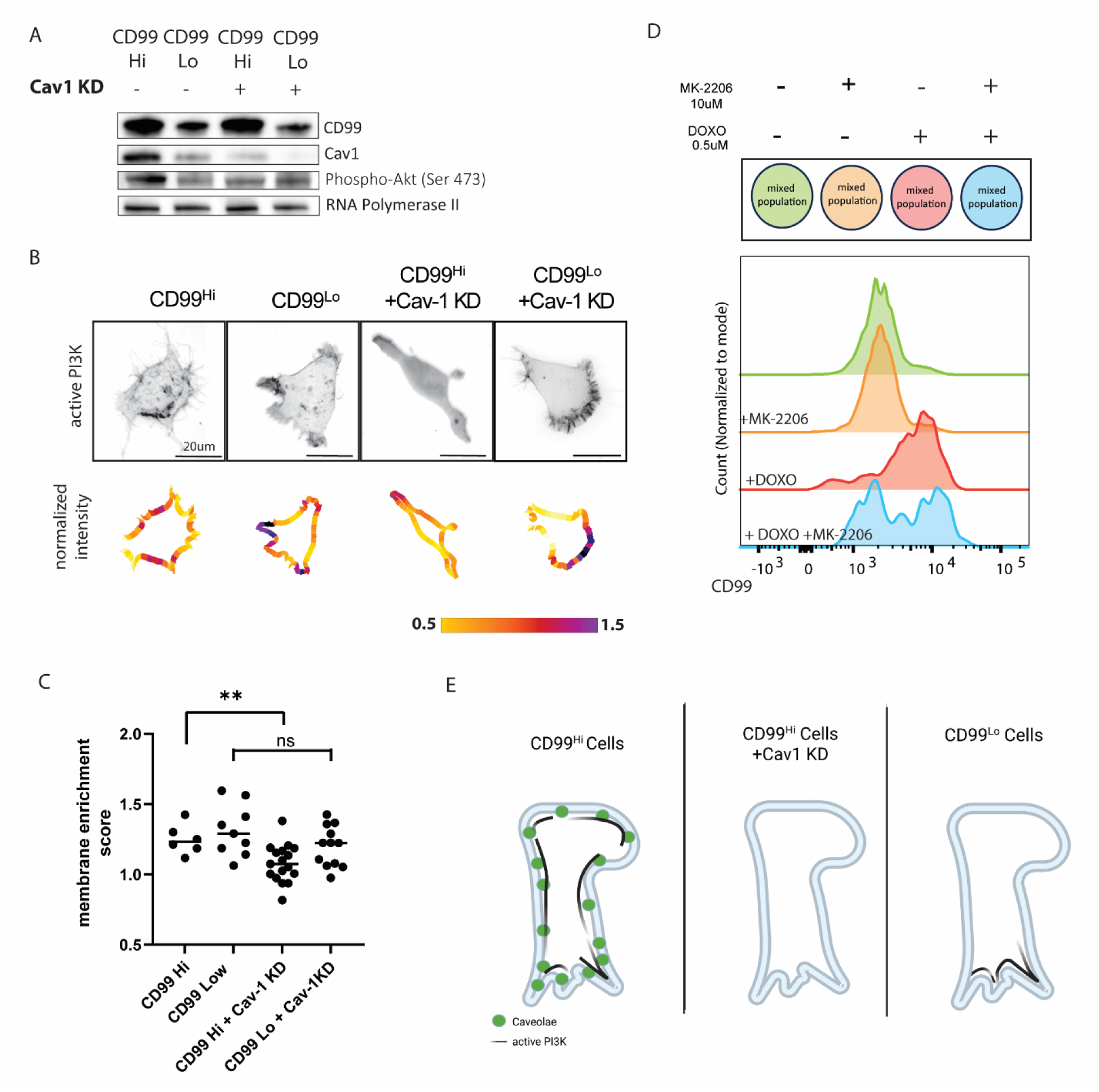
Caveolin-1 regulates PI3K/AKT signaling in CD99^Hi^ Cells. **(A)** Western blot depicting higher Cav1 expression and increased active AKT signaling in CD99^Hi^ compared to CD99^Lo^ cells, which is lost upon Cav-1 KD in CD99^Hi^ cells (n=2 independent repeats). **(B)** TC71 CD99^Hi^ and CD99^Lo^ cells, and CD99^Hi^ and CD99^Lo^ cells +Cav1 KD, expressing PI3K activation sensor AktPH-GFP (active PI3K). (Top) Maximum Intensity Projection, (Bottom) Measured membrane enrichment of active PI3K on cell boundary. Pseudocoloring is normalized to mean intensity of cell volume. **(C)** PI3K activity membrane enrichment score (n = 6, 9, 17, and 12 cells, for CD99^Hi^, CD99^Lo^ cells, CD99^Hi^ + Cav1 KD, and CD99^Lo^ cells +Cav1 KD respectively. (** p-value <0.005,KS test). Membrane enrichment score is reduced upon Cav-1 KD in CD99^Hi^ but not CD99^Lo^ cells. **(D)** Flow cytometric analysis of CD99 expression in mixed populations of TC71 cells following treatment with the AKT inhibitor MK-2206 (10 μM), doxorubicin (DOXO; 0.5 μM), or the combination of MK-2206 and DOXO for 48 h. DOXO treatment resulted in enrichment of the CD99^Hi^ population, whereas combined treatment with MK-2206 and DOXO abolished this enrichment, resulting in a distribution similar to untreated cells. Treatment with MK-2206 alone had minimal effect on CD99 distribution. **(E)** Summary of proposed mechanism. CD99^Hi^ cells express high levels of Cav-1 and contain abundant caveolae, which alter membrane properties to stabilize active PI3K at the cell surface and promote downstream signaling. In contrast, CD99^Hi^ cells with reduced Cav-1 expression lack caveolae, resulting in impaired recruitment of active PI3K to the membrane and lower signaling. CD99^Lo^ cells do not rely on Cav-1 expression and appear to regulate PI3K signaling by other means.

The downstream effects of PI3K/Akt signaling on diverse cell functions rely on recruitment of the active molecule to the cell surface^51^. We therefore applied a high-resolution imaging approach to measure the levels and organization of PI3K activity at the surfaces of individual TC71 cells. We employed a GFP-tagged Akt-PH construct as a reporter of the local concentration of PI3K-activated lipid products^52, 53^ in CD99^Hi^ cells, CD99^Lo^ cells, and CD99^Hi^ cells with Cav-1 KD, seeded on glass coverslips (Figure 5B). Active PI3K localization was quantified using a membrane-enrichment score, defined for each cell as the 90th percentile of local membrane fluorescence intensity normalized to the mean fluorescence intensity within the cell interior. The 90th percentile was used to capture regions of strong membrane enrichment while limiting sensitivity to extreme values and measurement noise. Both CD99^Hi^ and CD99^Lo^ cells showed comparable membrane enrichment scores of active PI3K (Figure 5B-C). Strikingly, Cav-1 knockdown markedly reduced active PI3K membrane enrichment score in CD99^Hi^ cells (Figure 5B-C, S4D). In contrast, Cav-1 knockdown had no marked effect on active PI3K enrichment in CD99^Lo^ cells. Control experiments using a lipophilic membrane dye showed no significant differences in membrane enrichment score between conditions (Figure S4B,C), indicating that the observed changes are specific to PI3K signaling. We also assessed active PI3K membrane enrichment in CD99^Hi^ NCH-EWS5 cells, with and without Cav-1 KD. Although Cav-1 KD was associated with a lower mean enrichment score, this difference did not reach statistical significance, potentially because of the high variability observed in cells with endogenous Cav-1 expression (Figure S4E).

To determine whether AKT signaling contributes functionally to the survival advantage of CD99^Hi^ cells, we examined the effects of pharmacological AKT inhibition in the context of chemotherapeutic stress (Figure 5D). Treatment with the AKT inhibitor (MK-2206,10 μM) alone had little effect on the relative abundance of CD99^Hi^ and CD99^Lo^ cells within a mixed population. However, while treatment with DOXO (0.5 μM) alone resulted in enrichment of the CD99^Hi^ population as previously observed, this enrichment was abolished by the combination of DOXO and MK-2206. Thus, inhibition of AKT phenocopied the effects of Cav-1 knockdown, indicating that AKT signaling is required for the survival advantage of CD99^Hi^ cells under chemotherapeutic stress. Together, these findings support a model in which Cav-1 selectively promotes survival of CD99^Hi^ cells through regulation of PI3K/AKT signaling (Figure 5E).

## DISCUSSION

In this study, we uncover a subpopulation of EwS cells characterized by distinctions in transcriptional signature, cell morphology, and growth *in vivo.* These cells can be identified and isolated through flow cytometry based on their high expression of the cell surface receptor CD99, a glycoprotein often used as a diagnostic marker in EwS^54^. Functional characterization of CD99^Hi^ cells demonstrates a survival advantage in certain tissues *in vivo* and Cav-1 dependent survival in response to a chemotherapeutic drug. Our findings implicate Cav-1 as a regulator of EwS cell survival through cell-state-dependent modulation of PI3K signaling.

Cell state heterogeneity is a major factor in driving drug resistance and metastasis in cancer ^1^, creating an urgent clinical need to better understand the underlying mechanisms. In EwS, as in many pediatric cancers^4^, mutational tumor burden beyond a single driver mutation is low, suggesting that non-genetic mechanisms are a primary factor in modulating cell state heterogeneity. Several previous studies have identified variability in expression levels of the oncogene EWSR1::FLI1 or its transcriptional signature as a driver of distinct EwS cell states ^8, 39, 43, 55^, largely through experimental knockdown approaches. Notably, both CD99 and Cav-1 are direct targets of EWSR1:FLI1^56^. However, the differences between CD99^Hi^ and CD99^Lo^ cells described in this work appear independent of EWSR1::FLI1 expression level. While CD99 served as a convenient marker of the distinct cell states, our results do not support a direct functional role for CD99 itself in mediating the phenotypes associated with the CD99^Hi^ state. Cav-1 depletion did not alter CD99 expression, and conversely, CD99 knockdown did not alter Cav-1 levels in CD99^Hi^ cells, indicating that CD99 expression and Cav-1-associated signaling are functionally separable. Rather, CD99 expression provided a robust means to prospectively isolate a subpopulation with distinct signaling, morphological, and survival phenotypes. CD99^Hi^ cells display increased neural differentiation, as well as survival advantages and repressed tumor growth *in vivo,* phenotypes which have previously been associated with reduced CD99 expression^54, 57^. Despite being routinely profiled in EwS^5^, endogenous heterogeneity in CD99 expression with functional consequences has not previously been described. The emergence of the CD99^Hi^ state only upon non-enzymatic passaging may explain why this subpopulation of cells was overlooked in past studies. Together, our findings therefore define a previously uncharacterized EwS cell state with distinct functional behavior, expanding our understanding of non-genetic heterogeneity in this tumor.

We pursued Cav-1 as a potential driver of distinct cell states in EwS cells due to its strong differential expression levels across three EwS cell lines with identified CD99^Hi^ and CD99^Lo^ cell populations, and marked enrichment of Cav-1-driven caveolae in the CD99^Hi^ cells in TC71 cells. Importantly, these experiments identify Cav-1-dependent membrane organization, rather than CD99 itself, as the mechanistic correlate of the CD99^Hi^ state. Depletion of Cav-1 altered activation of downstream signaling pathways, and the ability of cells to survive *in vivo* or under drug challenge. In contrast, CD99^Lo^ cells lacked detectable caveolae, exhibited distinct growth and survival phenotypes *in vitro* and *in vivo*, and show altered organization of downstream signaling pathways, suggesting that the two populations may rely on distinct Cav-1-associated mechanisms to regulate survival. Together, these findings suggest that the relationship between Cav-1 and signaling is cell-state dependent, although the underlying mechanisms and their generalizability beyond the TC71 model remain to be established.

Cav-1 is known to regulate several major signaling pathways, such as Src-family kinases, receptor tyrosine kinases, and RAS signaling^16, 19, 20, 20, 31, 32^. Its effects on PI3K/Akt survival signaling pathway appear to be context dependent, with evidence for both activation^31, 32, 34^ and inhibition^58–60^, although the underlying molecular mechanisms remain incompletely understood. In our system, CD99^Hi^ cells localized active PI3K to the plasma membrane to a similar degree as CD99^Lo^ cells, yet exhibited higher levels of downstream phospho-AKT, suggesting more efficient propagation of PI3K-activated lipid signaling to AKT activation. Both enhanced AKT signaling and membrane recruitment of active PI3K were dependent on Cav-1 in CD99^Hi^ cells but not CD99^Lo^ cells. Together, these findings support a model in which Cav-1-dependent membrane organization enhances the coupling of PI3K-derived lipid to downstream AKT activation. While Cav-1 knockdown also reduced survival and proliferation in CD99^Lo^ cells, the distinct enrichment of PI3K/AKT signaling observed in CD99^Hi^ cells suggests that Cav-1 may regulate survival through different downstream mechanisms across these populations. In addition, while we demonstrated that Cav-1 knockdown impaired formation of caveolae in CD99^Hi^ cells, we also consider the possibility that Cav-1 knockdown may exert caveolae-independent effects, as reported previously ^61, 62^.

PI3K/Akt signaling has also been implicated in Ewing sarcoma biology^63–65^. Evidence suggests that EWSR1::FLI1 transcriptionally reinforces PI3K/Akt pathway activity ^66^, while PI3K/Akt signaling positively regulates EWSR1::FLI1 expression ^67^, forming a potential feed-forward loop. Although we did not observe differences in EWSR1::FLI1 expression between the subpopulations identified in this study, it remains possible that EWSR1::FLI1 transcriptional activity differs between these states and could regulate Cav-1 dependent PI3K/Akt signaling. Determining whether this interaction occurs, as well as identifying the extrinsic cues that promote the CD99^Hi^ state and the membrane properties that support its Cav-1-dependent signaling, may reveal cell-state-specific vulnerabilities in EwS.

## METHODS

### Cell culture and cell line generation

Cells were cultured at 5% CO2 and 21% O2. TC71 and RDES cells were cultured using RPMI (Gibco) supplemented with 10% FBS and 1% antibiotic antimycotic. For enzymatically passaged cells, cells were treated with 0.05% Trypsin-EDTA (GIBCO 15400054) up to 5 minutes. For mechanical passaging, cells were grown in T125 cell-culture treated flasks (Corning 430825), and iteratively washed with PBS by tapping of flask to detach cells. Cells were then reseeded in the same flask for 2-6 weeks until mixed cell morphologies of the CD99^HI^ and CD99^LO^ cell states were readily observed.

TC71 Cav1, EF1, and CD99 shRNA lines cells were generated using the pZIP shRNA Promoter selection kit (https://www.transomic.com/cms/home.aspx). shRNA knockdown was induced by continuous 1 µg/ml tetracycline in cell culture, and shRNA knockdown induction was verified by RFP reporter expression and Western blotting. For each target gene, a single shRNA construct was used for this study.

Fluorescent constructs were introduced into TC71 cells using the pLVX lentiviral system (Clontech) and selected using antibiotic resistance to either puromycin or neomycin. The GFP-FTractin and mRuby2-FTractin constructs contain residues 9–52 of the enzyme IPTKA30 fused to eGFP or mRuby2, respectively. pLenti6-H2B-GFP was acquired from Addgene (generated by Fuchs lab, New York University, plasmid # 25999; Addgene). The GFP-AktPH construct was obtained from the laboratory of Jason Haugh (North Carolina State University, Raleigh NC)^68^ and cloned into the pLVX-IRES-puro vector (Clontech).

Endogenously tagged TC71 Cav-1-mClover was generated through CRISPR-based knock-in strategy. Briefly, two guides were generated using CRISPOR^69^ (gRNA1: GCATCCCGATGGCACTCATC and gRNA2: GACCCACTCTTTGAAGCTGT), and were cloned into plasmid pSpCas9(BB)-2A-Puro (Feng Zhang lab, plasmid # 62988, Addgene). The donor vector was generated using plasmid pMA-Tia1L which was a gift from Tilmann Bürckstümmer^70^.

PDX lines NCH-EWS1 and NCH-EWS5 were obtained from PDX core of Greehey Children’s Cancer Research Institute (UT Health San Antonio). For NCH-EWS5, a short term cell culture line was generated for NCH-EWS5 from subcutaneous xenografts in mice, enabling lentiviral infection of GFP-AktPh and Cav1 shRNA constructs for imaging experiments.

### RPPA

Functional Proteomics Reverse Phase Protein Array (RPPA) was performed by the RPPA core at MD Anderson Cancer Center. Cellular proteins were denatured in a 1% SDS + 2-mercaptoethanol buffer solution and diluted in five 2-fold serial dilutions in dilution lysis buffer. Serially diluted lysates were arrayed on nitrocellulose-coated slides (Grace Bio-Labs) by the Quanterix (Aushon) 2470 Arrayer (Quanterix Corporation). A total of 5808 spots were arrayed on each slide including spots corresponding to serially diluted (1) standard lysates, and (2) positive and negative controls prepared from mixed cell lysates or dilution buffer, respectively. Each slide was probed with a validated primary antibody plus a biotin-conjugated secondary antibody. Antibody validation for RPPA is described in the RPPA Core website: https://www.mdanderson.org/research/research-resources/core-facilities/functional-proteomicsrppacore/antibody-information-and-protocols.html. Signal detection was amplified using an Agilent GenPoint staining platform (Agilent Technologies) and visualized by DAB colorimetric reaction. The slides were scanned (Huron TissueScope, Huron Digital Pathology) and quantified using customized software (Array-Pro Analyzer, Media Cybernetics) to generate spot intensity.

Relative protein level for each sample was determined by RPPA SPACE (developed by MD Anderson Department of Bioinformatics and Computational Biology, https://bioinformatics.mdanderson.org/public-software/rppaspace/)^71^ by which each dilution curve was fitted with a logistic model. RPPA SPACE fits a single curve using all the samples (i.e., dilution series) on a slide with the signal intensity as the response variable and the dilution steps as the independent variable. The fitted curve is plotted with the signal intensities, both observed and fitted, on the y-axis and the log2 concentration of proteins on the x-axis for diagnostic purposes. The protein concentrations of each set of slides were then normalized for protein loading. Correction factor was calculated by (1) median-centering across samples of all antibody experiments; and (2) median-centering across antibodies for each sample. Results were then normalized across RPPA sets by replicates-based normalization as described^72^. Details of the RPPA platform as performed by the RPPA Core are described in^73^.

### Transcriptomics

For single cell RNA-seq, RDES and TC71 cell lines were dissociated in Tryp-LE express enzyme (GIBCO), resuspended in 0.04% BSA in PBS, and passed through a 70um filter before diluting to 1000 cells/uL. NCH-EWS1 and SJEWS001321_X1 PDX lines were kindly provided by Peter Houghton (University of Texas Health Sciences center San Antonio), and St. Jude Children’s Research Hospital Childhood Solid Tumor Network (SJEWS001321_X1), respectively. For PDXs, tumor chunks >5 grams were dissociated using Trypsin and Collagenase, treated with a combination of DNAse and Magnesium Chloride to dissolve DNA, and passed through a 40um filter and treated with a Red Blood Cell lysis buffer before resuspending in 0.04% BSA in PBS and diluting to 1000 cells/uL. Samples were processed as in 10× Genomics Chromium 3’ V3.1 with Cell Multiplexing kit and pooled together evenly for a final concentration of 1500 cells/µL in 0.04% BSA in PBS. Single-cell RNA-seq libraries were prepared using 10× Genomics Chromium 3′ V3.1 kit, following the standard protocol.

Single-cell RNA sequencing (scRNA-seq) data were generated for two samples: NCH-EWS1 PDX, and TC71 cell line using the 10x Genomics platform. The data were processed using the Cell Ranger pipeline (version 6.0.0) to generate gene-barcode matrices, which were subsequently analyzed in Seurat (version 5.0.2). For each sample, raw count matrices, barcodes, and feature files were loaded into Seurat to create individual Seurat objects. Quality control was meticulously performed to exclude cells with extreme total RNA counts (nCount_RNA) and detected features (nFeature_RNA), with thresholds set based on the 5th and 95th percentiles for RNA counts, and the 10th and 90th percentiles for detected features. Additionally, cells with high mitochondrial content, indicative of potential cell stress or apoptosis, were filtered out by excluding cells with mitochondrial transcript percentages exceeding 15%. Data normalization was conducted using the “LogNormalize” method with a scale factor of 10,000, and highly variable features were identified using the “vst” method with 2,000 features selected for downstream analysis. Principal component analysis (PCA) was applied for dimensionality reduction, followed by clustering using the Louvain algorithm and visualization using Uniform Manifold Approximation and Projection (UMAP). Cell cycle scoring was performed by classifying cells into G1, S, and G2/M phases based on established markers. Specifically, cells in the G1 phase were subsetted for more focused analysis, ensuring that cell cycle effects were minimized in the downstream analyses. Gene Set Enrichment Analysis (GSEA) was conducted on the differentially expressed genes, utilizing pathways relevant to the EWSR1::FLI1 fusion as described by Siligan and Torchia. Furthermore, marker gene expression, such as CD99, was profiled across the different clusters, and volcano plots were generated to highlight significant differentially expressed genes.

### Doxorubicin treatment assays

For acute chemotherapy challenge experiments, mixed populations of CD99^HI^ and CD99^LO^ TC71 cells were treated with doxorubicin (DOXO; 0.5 μM or 10 μM, as indicated) for 48 hours. Following treatment, cells were harvested and processed for flow cytometry as described above. Cell viability was assessed by Live/Dead staining (ThermoFisher L34973) and flow cytometric analysis. CD99^HI^ and CD99^LO^ populations were defined based on CD99 fluorescence intensity as described in the Flow Cytometry section. he relative abundance of surviving CD99^HI^ and CD99^LO^ cells was quantified among viable cells and compared with untreated controls using FlowJo software (BD Biosciences. For Cav-1 knockdown experiments, shRNA expression was induced with tetracycline 48hrs prior to doxorubicin treatment as described above.

### Flow Cytometry

For flow cytometry, TC71 cells were dissociated in Tryp-LE express enzyme, resuspended in 5% FBS in PBS, and passed through a 70um filter before diluting to 1X10^6 cells/mL. For live cell staining of CD99, cell suspension was incubated with mouse anti-human PE-CD99 antibody (BD Pharmingen 555689, 1:6). For shRNA expressing cell lines expressing RFP, cell suspension was incubated with rabbit monoclonal CD99 antibody (Abcam EPR3096 1:50) followed by secondary anti-rabbit antibody conjugated to AlexaFlour 405 (Abcam ab175652, 1:1000). For viability staining, cells were additionally stained with 1:1000 Live-Dead fixable Far-Red stain (ThermoFisher L34973) prior to flow cytometry. Samples were analyzed or sorted based on CD99 detected fluorophore intensity using BD FACSAria Zelda (BD BioSciences) flow cytometers. Flow cytometry plots were visualized using FloJo Software (BD Bioscences).

### Western Blots

Primary antibodies used for western blotting were anti-CD99 (Abcam; EPR3096), Fli1 (Abcam; ab133485), Tubulin, Cav-1 (Cell Signaling; #3267), pAKT (Cell Signaling; #4060), AKT (Cell Signaling; #2920), Histone H3 (Cell Signaling: #9715), Phospho-PI3K (Invitrogen, #17387), PI3K (Cell Signaling, #42925).

### Zebrafish husbandry and xenografts

*Danio rerio* were maintained in an Aquaneering aquatics facility according to industry standards. Vertebrate animal work was carried out by protocols approved by the institutional animal care and use committee at UT Southwestern, an Association for Assessment and Accreditation of Laboratory Animal Care accredited institution. Wild Indian Karyotype (WIK) wild-type lines used were obtained from the Zebrafish International Resource Center (zebrafish.org).

For xenografts of TC71-Ftractin-mRuby2 cells, cells were sorted for CD99^HI^ and CD99^LO^ populations, then expanded prior to injection. Preparation of cell mixture consisted of 90% mRuby2-Ftractin labeled cells and 10% unsorted H2B-GFP labeled cells, or 100% mRuby2-Ftractin labeled cells, as indicated. Cell mixtures (2 × 10^6^ cells) were run through a 70-µm cell strainer and resuspended in 5% FBS in PBS mixture for optimal cell viability.

Zebrafish embryos were collected, dechorionated, and treated with 0.1 mM phenylthiourea starting at 24 h post fertilization to prevent pigmentation. Xenografts were generated by microinjection of cell mixture into zebrafish larvae (2 days post fertilization [dpf]) with glass capillary needles into HBV. For each cell xenograft group, cells were injected into 3-aminobenzoic acid ethyl ester (Tricaine)–anesthetized embryos. Injected zebrafish larvae were incubated for 1–2 d in 34°C before imaging.

For immunostaining of zebrafish larvae, anesthetized embryos were fixed in 4% paraformaldehyde/1XPBS for 2 h in scintillation vials, rinsed in PBS +0.5% Tween, and stored in PBS up to 2 wk. Immunostaining was carried out as previously described (Miller et al., 2015). Primary antibodies were rabbit anti-GFP (ab290; Abcam) and mouse anti-Phospho-Histone H3 (9706S; Cell Signaling). TUNEL assay was performed using the ApopTag Green In-Situ Apoptosis Detection Kit (Millipore).

### Mouse xenografts

All experiments involving animals were reviewed, approved, and monitored for compliance by the UT Southwestern Institutional Animal Care and Use Committee.

Lenti-luciferase-P2A-Neo was acquired from Addgene (Vakoc lab, Cold Spring Harbor Laboratory, plasmid: #105621; Addgene), and introduced into TC71 cells, which were subsequently sorted for *CD99 High and CD99 Low* populations and expanded prior to xenografts. Maintenance of isolated cell states was validated by distinct cell morphology in cell culture. Orthotopic intratibial xenografts were performed in NOD *scid* gamma chain knockout (NSG) mice, following a previously established protocol^74^. Briefly, the top of the right tibia was punctured with a 25 gauge needle, followed by injection of 5×10^5^ cells suspended in 5ul media into the medullar cavity of the bone using a 27 gauge needle from a tuberculine syringe (Cat 26040, Exelint, Int. Co, Redondo Beach, CA) coupled to a 0.1 ml Hamilton syringe (Cat 8001, Hamilton Co, Reno, NV).

Functional imaging to monitor tumor growth in live mice was performed using the IVIS Spectrum System (PerkinElmer). At each imaging session, 80 µL of a Luciferin working solution (40 mg/mL in PBS without calcium or magnesium) was administered. Mice were imaged at 0, 7, 14, 21, 28, 35, and 42 days post-injection. Regions of interest (ROIs) around the injection site were manually delineated for quantitative analysis of total flux [photons/second]. Luciferin signals observed in the liver were excluded from the analysis. At the end of the experiment, tumor-bearing Grossly trimmed tibia samples and their encapsulating musculature and non-tumor bearing counter-parts from the same mouse were harvested for further evaluation.

For micro-Computer Tomography (microCT) scans, formaline-fixed samples were scanned using a Scanco Medical μCT 35. Tibias were scanned at an isotropic voxel size of 3.5μm, with peak tube voltage of 55 kV and current of 0.145 mA (μCT 35; Scanco). A Gaussian filter (s = 0.8) with a limited, finite filter support of one was used to suppress noise. Mineralized bone was segmented from air and soft tissues by intensity thresholding. Scanco Medical software was used to determine bone volume.

### Mouse tumor histology

Grossly trimmed tibia samples and their encapsulating musculature from each condition were formalin-fixed, decalcified 14% ethylenediaminetetraacetic acid disodium salt dihydrate in paraffin-embedded, and sectioned at 4 µm thickness. Serial sections were used for Hematoxylin and Eosin staining (H&E), immunohistochemistry (IHC), immunofluorescence (IF), and TUNEL assay. All histology samples were evaluated by an expert pathologist.

Confirmation of proliferating cells and cell death were assessed by IF of Ki-67 (Leica, Cat# KI67-MM1-L-CE, 1:300) and DeadEnd Fluorometric TUNEL assay (Promega; Cat #: G3250), respectively, and counterstained with propidium iodide. IHC of CD99 staining was done with rabbit monoclonal CD99 antibody (Abcam EPR3096 1:50), using the SignalStain Boost IHC detection reagent (Cell Signaling Cat #8114), as directed.

All slides were digitally scanned on a Nanozoomer S60 (Hamamatsu) or Axioscan 7 (Zeiss) systems. Measurements of area were made through manual annotations of ROIs on images using NDP.view2 (Hamamatsu).

### Electron Microscopy

For each treatment, duplicate cell monolayers were grown on MatTek dishes. After an initial wash with PBS, the monolayers were washed with 0.1 M sodium cacodylate and then with 2.5% (v/v) glutaraldehyde in 0.1M sodium cacodylate buffer. After fixation, the monolayers were rinsed five times in 0.1 M sodium cacodylate buffer, then were post-fixed in 1% osmium tetroxide and 0.8 % K3[Fe(CN)6] in 0.1 M sodium cacodylate buffer for 1 hour at 4 degrees C. Cells were rinsed with water and en bloc stained with 2% aqueous uranyl acetate overnight at 4 degrees C. After five rinses with water, specimens were dehydrated with increasing concentration of ethanol at 4 degrees C, infiltrated with Embed-812 resin and polymerized in a 60°C oven overnight. Embed-812 disc was removed from MatTek plastic housing by submerging the dish in liquid nitrogen. Two discs were sandwiched together side to cell side by putting a drop of Embed-812 and polymerized in a 60°C oven overnight. Sandwiched discs were sectioned perpendicular with a diamond knife (Diatome) on a Leica Ultracut UCT (7) ultramicrotome (Leica Microsystems) and collected onto copper grids, post stained with 2% Uranyl acetate in water and lead citrate. TEM images were acquired on a JEOL-1400 Plus transmission electron microscope equipped with a LaB6 source operated at 120 kV using an AMT-BioSprint 16M CCD camera.

### Imaging

Light-sheet fluorescence microscopes were used to image the zebrafish larvae within a custom-made mount. Fixed zebrafish larvae were embedded in 1% low-melting agarose into sonicated, ethanol-washed fluorinated ethylene propylene tubes (Zeus, HS 029EXP/.018 REC). In this case, the tube is held by a custom-made mount, which allows 3D translation and rotation in our light-sheet microscopes. A conventional mSPIM^75^ was used as previously described^41^. Briefly, a 40× NA 0.8 water-dipping objective (CFI Apo NIR 40XW; Nikon Instruments) was used for detection, at room temperature in PBS. A 200-mm focal length tube lens (58-520; Edmund Optics) that formed the image on an sCMOS camera (ORCA-Flash4.0; Hamamatsu Photonics). These conditions provide a resolution of 380 nm laterally and about 1.4 µm axially. A 10× NA 0.28 objective (M Plan Apo 10×, 378-803-3; Mitutoyo) was used for light-sheet illumination. The thickness and the length of the light-sheet were controlled by a variable slit (VA100C; Thorlabs) placed at the conjugate plane of the back focal plane of the illumination objective.

For live imaging on glass, F-Tractin-GFP and GFP-AktPH cells were mounted on a glass-bottom 35 mm cover dish (MatTek). GFP-AktPH cells were stimulated with Insulin Growth Factor (IGF) prior to imaging. IGF stimulation was done using 1nM Human IGF-1 recombinant protein (Peprotech) for 15 minutes, following 4 hours serum starvation. In some cases, GFP-AktPH cells were additionally labeled with a lipophilic membrane dye (CellBrite Steady Membrane Dye, biotium, 1:1000) immediately prior to imaging. Mounted cells were subsequently imaged using a custom-built Oblique Plane Microscope (OPM), as previously described^76^. Briefly, a tilted light sheet was launched after a primary objective (25x NA 1.1, Nikon MRD77220) to illuminate the cells in a glass-bottom dish; the fluorescence was collected by the same objective and formed the image on the camera after passing through the optical train with the magnification of around 43.3. By scanning the light sheet using Galvo mirrors in the optical train, we can rapidly acquire the 3D image of the cells without any mechanical movement of either sample or objective. Also, using a single objective for both illumination and fluorescence collection frees the whole sample space above the objective and makes the system compatible with the standard sample mounting methods. The resolution of the system across the extended FOV was around 0.39, 0.43, and 1.22 µm in the x, y, and z directions, respectively.

To image dynamics of Cav-1 puncta, TC71 Cav1-mClover cells were imaged on a SoRA spinning disc confocal microscope (Nikon) at 100X magnification, NA1.45, 200ms exposure, at 2 second time intervals. PTRF and Cav1-mClover localization was imaged on LSM780 scanning confocal microscope (Zeiss) at 40X magnification, NA1.4.

### Image Analysis

For quantification of Cav-1 puncta dynamics, particle detection and tracking were performed using u-track (version 2.4; https://github.com/DanuserLab/u-track)^77^. The detection algorithm used was Gaussian-mixture model fitting, using default parameters. The mean localization precision per movie was 8.5-12.1 nm. The resulting detections were then tracked over time, using default tracking parameters. The detection and tracking results were verified using visual inspection as well as using the performance diagnostics included in u-track. Tracks lasting for at least 10 frames were then analyzed to obtain each track’s diffusion coefficient from its frame to frame displacements^78, 79^.

For 3D morphology classifications in zebrafish, images were segmented using the previously established 3D morphology pipeline^80^ to segment cell volumes and calculate local membrane curvature from single cell images. For the segmentation process, we smoothed the deconvoluted images with a 3D Gaussian kernel (σ = 3 pixels) and applied a gamma correction (ϒ = 0.7), then employed an Otsu threshold, followed by a 3D grayscale flood-fill operation to fill possible holes. The binary images of all cells were visually inspected for segmentation artifacts. Miss-segmented cells were excluded from the data set. Aspect Ratio was calculated on segmented volumes using “regionprops3” in MATLAB. Aspect ratio was defined as the ratio of the major to minor axis lengths of the fitted ellipsoid and was used as a quantitative measure of cell elongation, with larger values indicating more elongated morphologies and values approaching 1 indicating more rounded cells. 3D rendering were generated using ChimeraX^81^.

To quantify differences in tissue morphology between CD99^Hi^ and CD99^Lo^ cells xenografted in mice, we compared various aspects of nuclear morphology in Hematoxylin and Eosin (H&E) stained histological slides. First, tumor regions to be analyzed were manually annotated using QuPath^82^ on tissue images from 3 slides in each experimental group, scanned at 40X magnification (∼0.25 microns/pixel). Second, nuclei in these regions were segmented using the PanNuke version of the HoverNet model as implemented in the TIAToolbox^83^. Third, these images were downsampled to 20X magnification and their color intensities we normalized using the Macenko algorithm^84^ against a single (randomly selected) reference slide. Finally, we extracted 37 morphological features relating to size, shape, color, and texture of each individual nucleus, as previously described^85^. Then for each annotated tumor region we randomly selected neighborhoods of 25 nuclei: the number of neighborhoods was proportional to the area of the region but capped at 500 neighborhoods. Each neighborhood was then represented by a feature vector of the median value of individual features across constituent cells, which were then pooled across slides, z-score normalized, and visualized using T-distributed Stochastic Neighbor Embedding (t-SNE).

For analysis of active PI3K organization or membrane dye enrichment on the cell surface in 2D maximum intensity projection images, we segmented each cell and windowed its peripheral boundary. Each window was ≈ 2.88 µm x 2.88 µm. To window, we implemented the conformalized mean curvature flow deformation from the u-Unwrap3D framework^86^ (https://github.com/DanuserLab/u-unwrap3D) into the 2D u-Unwrap framework^87^ (https://github.com/DanuserLab/u-unwrap) to deform the 2D cell boundary, equipartitioned the cell boundary and grouped together the corresponding deformed points of each boundary partition. For intensity measurement, we measured the mean fluorescence signal within each window, and computed the ratio enrichment with respect to the mean fluorescence signal averaged over the whole cell area. To compare the intensity distribution shift across conditions using a single summary statistic we computed the 90th percentile of the mean window normalized intensity for each cell.

Plots and statistical tests were visualized and preformed using GraphPad Prism.

## ACKNOWLEDGMENTS

We would like to thank UT Southwestern Whole Brain Microscopy Facility (grant award number 1S10OD032267-01 to Denise Ramirez), Flow Cytometry Core Facility, Histo Pathology Core Facility, and Quantitative Light Microscopy Core Facility (grant award number 1S10OD028630-01 to Katherine Luby-Phelps), for their contributions to this work.

Funding for this work in the Danuser, Fiolka, and Amatruda labs has been provided by the grant 1U54CA268072 (NCI). Some microscope development in the Fiolka lab has been supported by R33 CA235254 (National Cancer Institute). D. Segal acknowledges funding from European Molecular Biology Organization Long-term Research Fellowship (ALTF 626-2018) and the grant K99 CA270285 (NCI). X. Wang and N.S. Williams acknowledge funding from the NCI (1U54CA231649) and institutional support from UTSW for the Preclinical Pharmacology Core. Funding in the Jaqaman lab is provided through R35 GM119619 (National Institute of General Medical Sciences) and the UT Southwestern Endowed Scholars program.

## DECLARATION OF INTERESTS

G.D. is member of the board of Glencoe Software, Inc. and an advisor for the Allen Institute of Cell Science. The authors declare no other competing interests.

## Notes

### Competing Interest Statement

G.D. is member of the board of Glencoe Software, Inc. and an advisor for the Allen Institute of Cell Science. As of August 2025, G.D. has been co-director of the Institute of Human Biology at Hoffmann La Roche in Basel. The authors declare no other competing interests.

### Summary of Updates

This manuscript has been revised substantially following review. Figures 2,3,4, and 5 have been revised, and writing has been reorganized to add clarity.

