## Supplemental data for "Caveolin-1-dependent signaling and survival in distinct Ewing sarcoma cell states"

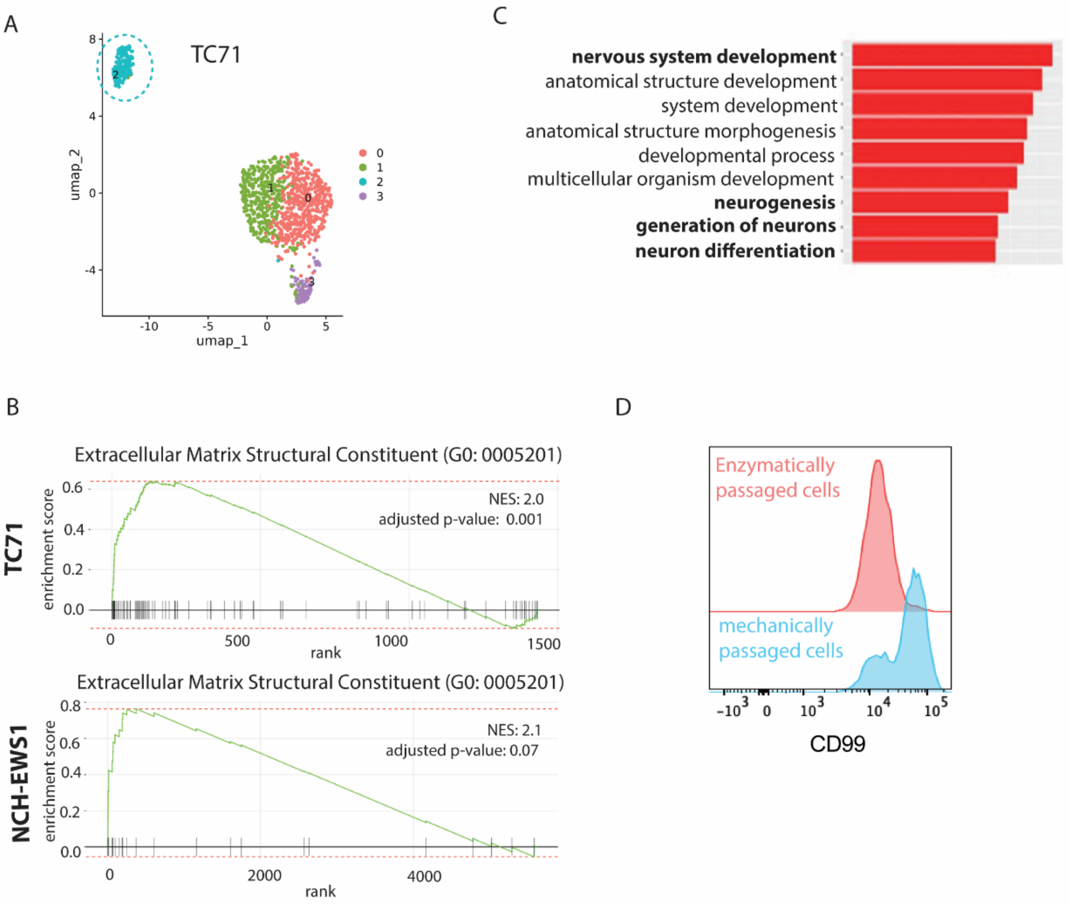


**Figure S1 (related to Figure 1): High levels of CD99 gene expression marks a distinct Ewing Sarcoma cell state. (A)** UMAP visualization of TC71 cells analyzed by scRNA-seq, with cluster numbers annotated. We identified a cluster (Cluster 2, highlighted) with a distinct transcriptional profile. **(B)** Gene Set Enrichment Analysis (GSEA) plot of Cluster 2 vs. all other clusters shows enrichment in $CD99^{Hi}$ cluster for an Extracellular Matrix Structural Constituent gene signature in TC71 (top) and NCH-EWS1 patient-derived xenograft (PDX) (bottom) cells (NES-normalized enrichment score). **(C)** Gene Ontology terms for $CD99^{Hi}$ cells show enrichment of neurally differentiated gene signatures relative to $CD99^{Lo}$ cells. **(D)** Flow Cytometric analysis showing bimodal distribution of CD99 expression in TC71 cells when passaged through mechanical passaging (blue) but not enzymatic passaging (red), highlighting that the $CD99^{Hi}$ population emerges only under certain culturing conditions.

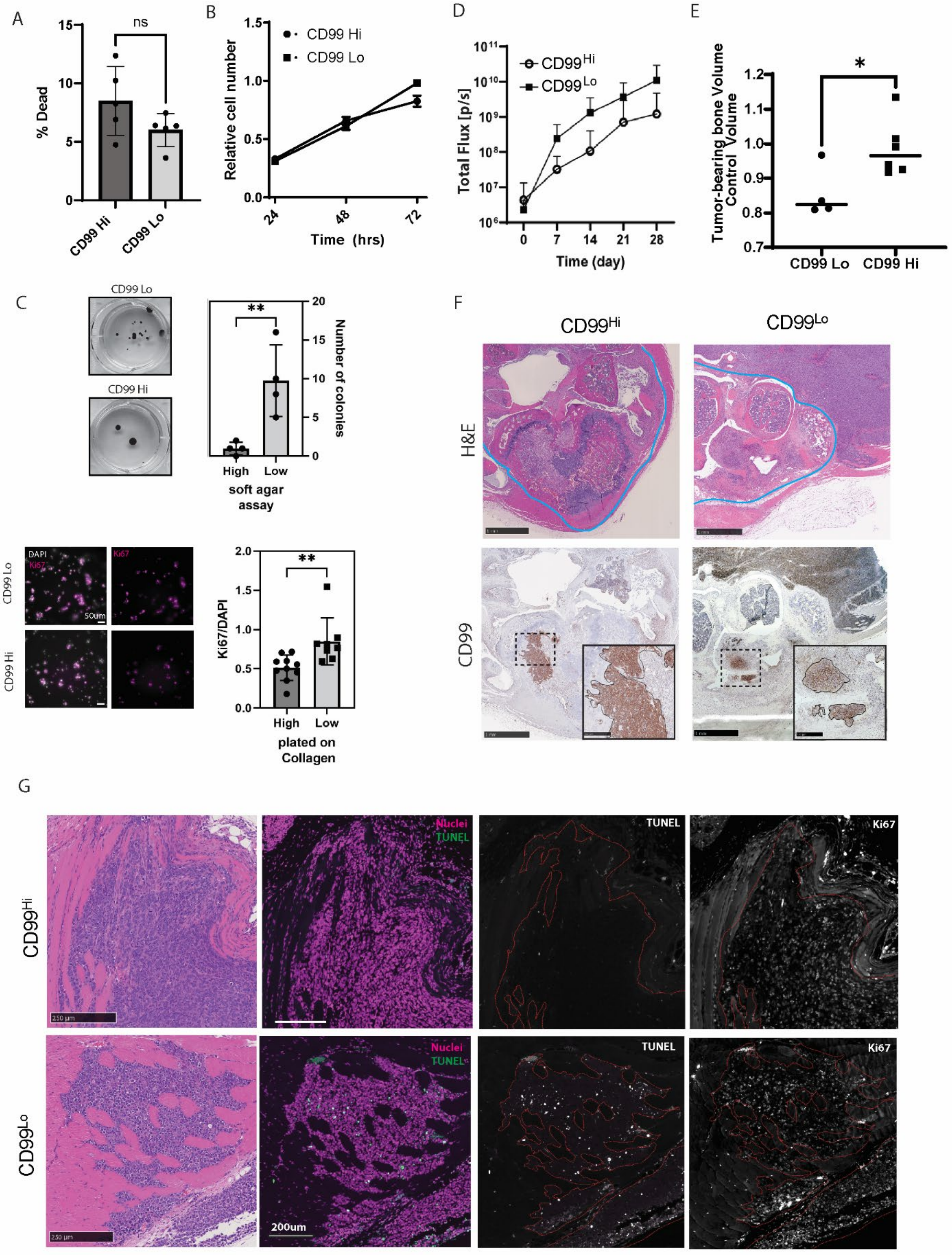

A
ns
% Dead
CD99 Hi
CD99 Lo
B
CD99 Hi
CD99 Lo
Relative cell number
Time (hrs)
D
CD99Hi
CD99Lo
Total Flux [p/s]
Time (day)
E
Tumor-bearing bone Volume
Control Volume
CD99 Lo
CD99 Hi
C
CD99 Lo
CD99 Hi
Number of colonies
High
Low
soft agar assay
DAPI
Ki67
50um
Ki67/DAPI
plated on Collagen
F
CD99Hi
CD99Lo
H&E
CD99
G
Nuclei
TUNEL
Ki67
250 µm
200um

**Figure S2 (related to Figure 3): CD99$^{Hi}$ and CD99$^{Lo}$ cells display differences in morphology, gene expression, and survival. (A-B)** No differences are observed in cell survival, measured by integration of Ethidium Homodimer (**A;** n = 5 independent repeats) or proliferation, measured by a cell viability assay (**B;** n = 3 independent repeats) in TC71 CD99$^{Hi}$ or CD99$^{Lo}$ cells grown in adherent culture conditions. **(C)** Cells plated on Collagen and stained for proliferation marker Ki67 (top, n=10 independent repeats) or soft agar assays (bottom, n=4 independent repeats) show significantly slower growth of CD99$^{Hi}$ cells in soft environments (**, $p<0.005$, t-test). **(D)** Total bioluminescent flux of TC71 in CD99$^{Hi}$ or CD99$^{Lo}$ luciferase-expressing cells in intratibial xenografts of immunocompromised mice, measured over time. Difference between curves is statistically significant (n = 6 mice, $p<0.05$, mixed-effect analysis). **(E)** Measurements in computer tomography scans of the bone volume of the tumor-bearing leg normalized by the volume of the non-tumor-bearing leg. Note that legs xenografted with CD99$^{Hi}$ cells (n = 6) display higher normalized volume compared to those xenografted with CD99$^{Lo}$ cells (n=4), suggesting reduced bony destruction in CD99$^{Hi}$ cells ($p<0.05$, t-test). **(F)** Representative histological sections of intratibial xenografts, showing tumor-bearing areas of either CD99$^{Hi}$ or CD99$^{Lo}$ cells, labeled with H&E (top), or chemically immunostained for CD99 (bottom, inset outlines CD99+ cells within enlarged region marked with dashed line). Boundary marking medullary cavity (med. cav.) shown with cyan line on H&E slide. Note the relatively large CD99+ tumor outside of the med. cav. and in the surrounding skeletal muscle in the CD99$^{Lo}$ tumor, with smaller tumor masses within the med. cav. In contrast, the CD99$^{Hi}$ tumor is present only in a relatively small area within the med. cav. **(G)** Histological sections of intratibial xenografts in immunocompromised mice, showing tumor-bearing areas outside the med. cav. of either CD99$^{Hi}$ or CD99$^{Lo}$ cells, labeled with H&E, and fluorescently stained for the apoptotic marker TUNEL or the proliferative marker Ki67, and with a propidium iodide nuclear counter stain, in serial sections. Red dashed outlines indicate tumor-bearing areas. Note the decreased TUNEL staining observed in the CD99$^{Hi}$ tumor compared to CD99$^{Lo}$ tumor.

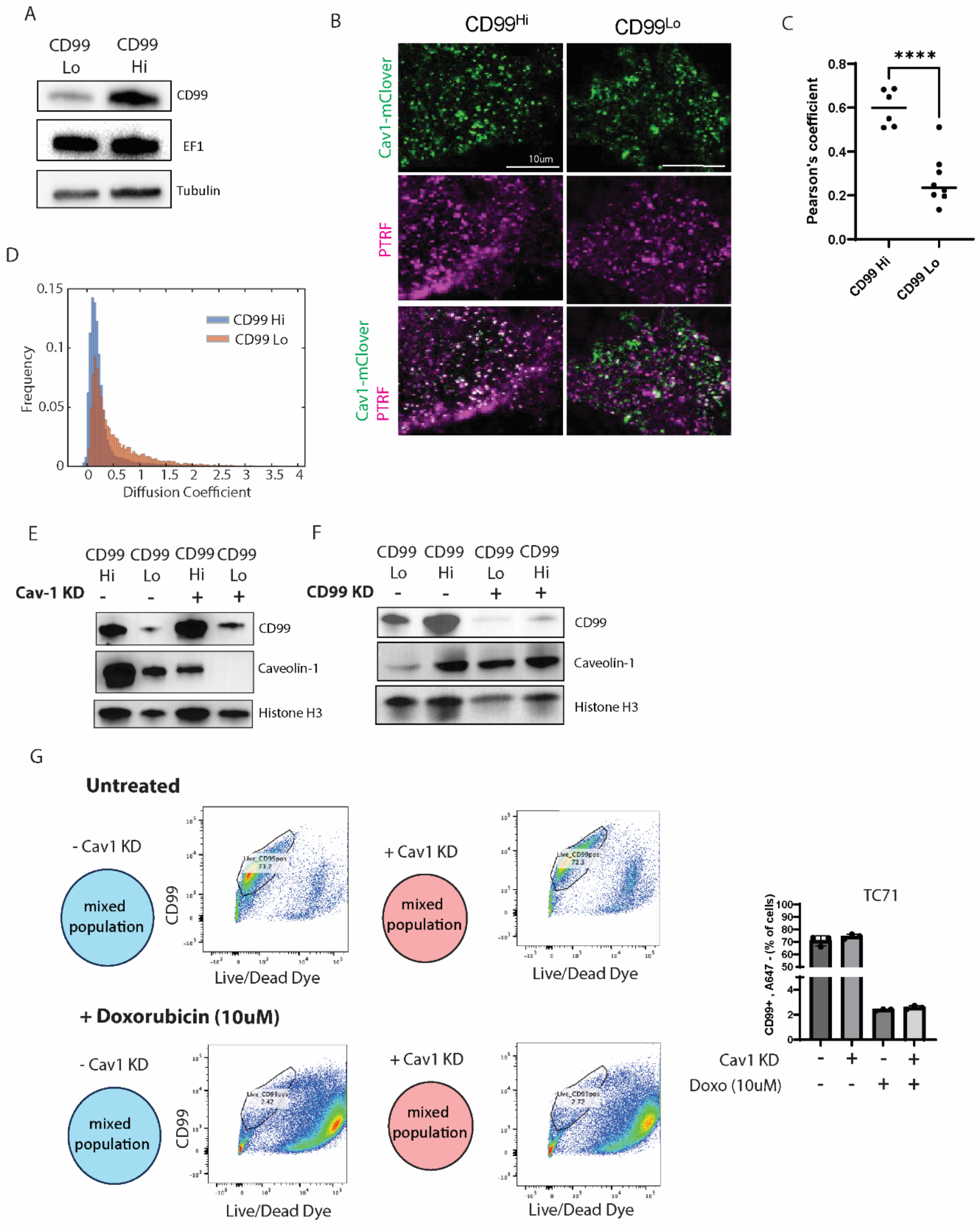
A
CD99 Lo
CD99 Hi
CD99
EF1
Tubulin
B
CD99Hi
CD99Lo
Cav1-mClover
10um
PTRF
Cav1-mClover PTRF
C
****
Pearson's coefficient
0.8
0.6
0.4
0.2
0.0
CD99 Hi
CD99 Lo
D
0.15
0.1
0.05
0
Frequency
CD99 Hi
CD99 Lo
0 0.5 1 1.5 2 2.5 3 3.5 4
Diffusion Coefficient
E
CD99 Hi
CD99 Lo
CD99 Hi
CD99 Lo
Cav-1 KD - - + +
CD99
Caveolin-1
Histone H3
F
CD99 Lo
CD99 Hi
CD99 Lo
CD99 Hi
CD99 KD - - + +
CD99
Caveolin-1
Histone H3
G
Untreated
- Cav1 KD
mixed population
CD99
Live/Dead Dye
+ Cav1 KD
mixed population
Live/Dead Dye
+ Doxorubicin (10uM)
- Cav1 KD
mixed population
CD99
Live/Dead Dye
+ Cav1 KD
mixed population
Live/Dead Dye
TC71
CD99+ , A647 - (% of cells)
100
90
80
70
60
50
4
2
0
Cav1 KD - + - +
Doxo (10uM) - - + +

**Figure S3 (related to Figure 4): Caveolin-1 emerges as a potential driver of CD99$^{Hi}$ State.** **(A)** Western blot depicting differences in CD99 expression but not EWSR1::FLI1 (EF1) expression between CD99$^{Hi}$ or CD99$^{Lo}$ TC71 cells (n=2 independent repeats). **(B-C)** Representative images (B), and quantification of colocalization by Pearson's coefficient (C) of TC71 CD99$^{Hi}$ or CD99$^{Lo}$ cells with endogenous expression for Cav1-mClover, and immunostained for PTRF show colocalization of Cav-1 and PTRF in CD99$^{Hi}$ but not CD99$^{Lo}$ cells (n=6,8 cells, respectively, t-test, ****p- value<0.0001). **(D)** Histogram of diffusion coefficient of individual caveolae, calculated from live imaging of endogenously tagged TC71 Cav1-mClover in CD99$^{Hi}$ (n = 10,063 puncta in 3 cells) and CD99$^{Lo}$ cells (n = 6,350 puncta in 12 cells) at 3s time intervals. Note higher diffusion of Cav-1 in CD99$^{Lo}$ cells. **(E)** Western blot showing efficiency of inducible shRNA knockdown of Cav-1, with no effect on CD99 expression upon reduced Cav-1 expression (n=3 independent repeats). **(F)** Western blot showing that inducible CD99 knockdown drives higher Cav-1 expression in CD99$^{Lo}$ cells but has little to no effect on Cav-1 expression in CD99$^{Hi}$ cells (n=3 independent repeats). **(G)** (left, middle) Flow cytometry-based viability assay of a mixed population of CD99$^{Hi}$ or CD99$^{Lo}$ cells, with (magenta) or without (blue) Cav-1 KD, untreated (top) or grown with Doxorubicin (10uM, bottom). The x-axis shows measurement of the A647-Live/Dead viability dye, while the y-axis shows measured intensity of CD99. The CD99+, A647 negative population is outlined. (right) Overall survival of cells, as detected by CD99+, A647- population with or without drug treatment and with or without Cav-1 KD.

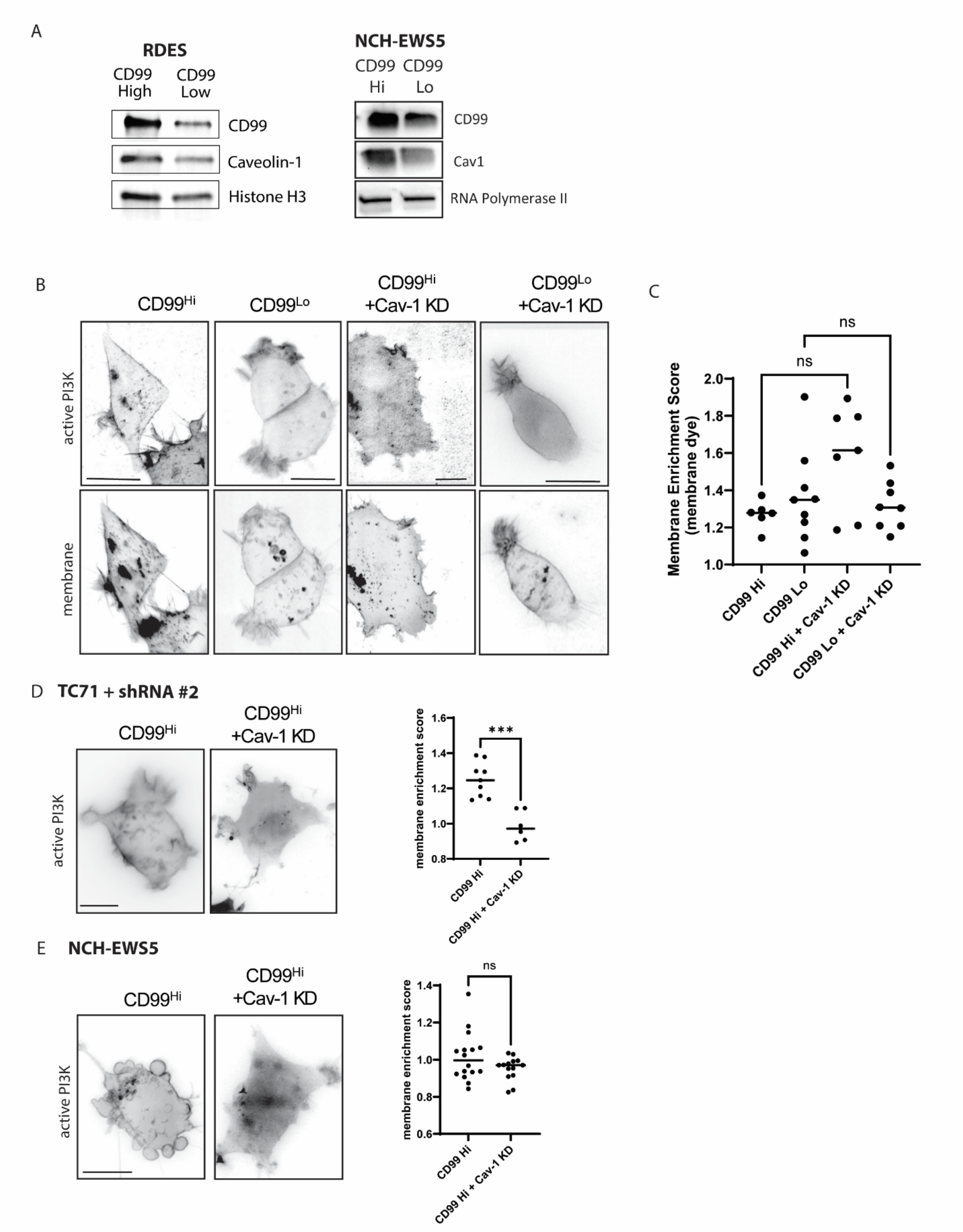


**Figure S4 (related to Figure 5): Cav-1 regulates PI3K/AKT signaling in CD99$^{Hi}$ cells (A)** Western blot depicting higher CD99 and Cav-1 expression in CD99$^{Hi}$ cells in two additional

Ewing Sarcoma cell lines: RD-ES (n=2), and NCH-EWS5 cells, a short-term culture derived from a patient-derived xenograft (PDX) (n=2). **(B)** Representative TC71 $CD99^{Hi}$ and $CD99^{Lo}$ cells, and $CD99^{Hi}$ and $CD99^{Lo}$ cells +Cav1 KD, expressing PI3K activation sensor AktPH-GFP (active PI3K, top), and stained with a lipophilic membrane dye (bottom). Scale bars = 20μm. **(C)** Membrane Enrichment Score of Membrane Dye on cell surface normalized to mean intensity of cell volume (n = 6, 9, 7, and 8 cells, for $CD99^{Hi}$, $CD99^{Lo}$ cells, $CD99^{Hi}$ + Cav-1 KD, and $CD99^{Lo}$ cells +Cav1 KD respectively (no significant differences, KS test). **(D)** Representative images (left) and quantification (right) of membrane enrichment of active PI3K in TC71 $CD99^{Hi}$ cells alone, or expressing a second shRNA construct driving Cav-1 KD. $CD99^{Hi}$ cells show reduced active PI3K upon Cav-1 KD (n=9 cells, 6 cells, respectively). Scale bars = 10μm, *** p-value <0.001, t-test. **(E)** Representative images (left) and quantification (right) of NCH-EWS5 $CD99^{Hi}$ cells, with or without Cav-1 KD showing membrane enrichment of active PI3K. While $CD99^{Hi}$ cells show higher variability in membrane enrichment score when expressing endogenous Cav-1 levels, no significant difference was detected upon Cav-1 KD (n = 16 cells, 14 cells, respectively. p = 0.10, t-test).

**Table S1 (separate excel file): Several Receptor Tyrosine Kinases are enriched in CD99 High Cells.** Results from a Functional Proteomics Reverse Phase Protein Array (RPPA) show differential expression in several key signaling proteins in $CD99^{Hi}$ Cells.
